# Multiple mechanisms mediate the suppression of motion vision during escape maneuvers in flying *Drosophila*

**DOI:** 10.1101/2022.02.03.478949

**Authors:** Philippe J. Fischer, Bettina Schnell

## Abstract

During voluntary behaviors, animals need to disable any reflexes that could interfere with the intended movements. With the optomotor response, for example, flies stabilize a straight flight path by correcting for unintended deviations sensed as panoramic motion of the surround. HS cells of the fly are thought to mediate optomotor responses to horizontal motion. During spontaneous flight turns, an efference copy acts on HS cells with the right sign to counteract the visual input elicited by the fly’s own behavior. Here, we investigated HS cell activity during looming-elicited turns in flying *Drosophila*. We show that looming stimuli themselves can influence the processing of panoramic motion stimuli in HS cells and that in addition, an inhibitory efference copy suppresses excitatory motion responses during turns, but only in a subset of HS cells. In conclusion, our findings support the notion that processing of sensory information is finely tuned to behavioral context.

## INTRODUCTION

Animals must be able to discriminate self-generated (reafferent) from external (exafferent) sensory input. While seminal work by von Holst and Mittelstaedt suggested that this can be achieved via an efference copy of the intended behavior that counteracts reafferent sensory input (Cullen, 2004; Von Holst and Mittelstaedt, 1950), its neuronal underpinnings have largely remained elusive.

In the optomotor response, for example, flies compensate for unintended deviations from a straight flight path, which they sense as motion of the surround or optic flow (Götz, 1968; Leonte et al., 2021; Mronz and Lehmann, 2008). However, this reflex-like compensation needs to be suppressed, when a fly performs a voluntary turn, which is also called a saccade. Saccades are rapid whole-body maneuvers that flies perform to change direction during flight (Muijres et al., 2015, 2014; Schilstra and Hateren, 1999; Schilstra and van Hateren, 1998). Saccades do not only occur spontaneously, but can be triggered by looming stimuli, indicative of an impending collision or approaching predator (Bender and Dickinson, 2006; Heisenberg and Wolf, 1979; Tammero and Dickinson, 2002). In head-fixed flight, intended saccades can be measured as fast changes in the difference between the left and right wing stroke amplitude (L-R WSA), which allows for studying their influence on visual processing in the fly brain (Heisenberg and Wolf, 1979; Maimon et al., 2010; Schnell et al., 2017). Saccades have been shown to follow a different motor program when executed spontaneously compared to when they are evoked by a looming stimulus (Dickinson and Muijres, 2016), but are indistinguishable in tethered flight quantified by changes in L-R WSA.

HS cells are a class of large-field lobula plate tangential cells that respond to horizontal motion in a directionally selective fashion (Hausen, 1982a, 1982b; Schnell et al., 2010). They receive ipsilateral input from columnar motion detecting cells on their dendritic trees, but also contralateral input from heterolateral lobula plate tangential cells (Haag and Borst, 2001). There are three HS cells per hemisphere, HSN, HSE and HSS, which differ in the position of their dendritic trees in the lobula plate and thus their receptive fields within the fly’s visual field (Hausen, 1982b; Schnell et al., 2010; Scott et al., 2002). Based on their response properties as well as activation and silencing experiments, they are thought to control stabilizing optomotor turns of the head and body during walking and flight (Busch et al., 2018; Fujiwara et al., 2017; Haikala et al., 2013; Kim et al., 2017), but have also been implicated in the processing of translational optic flow (Boeddeker et al., 2005). It has been shown that during spontaneous saccades an efference copy acts on HS cells, which has the appropriate sign to counteract responses to the visual stimulus that would be caused by the flies’ self-motion (Kim et al., 2017, 2015). However, it remained largely unclear how this effect influences processing of large-field motion in HS cells, which are thought to underlie stabilizing optomotor responses to horizontal motion.

We studied how saccades influence visual processing in HS cells. To be able to time the execution of saccades, we used looming stimuli to trigger them and combined them with presentation of panoramic horizontal motion. Using whole-cell patch-clamp recordings during head-fixed flight (Joesch et al., 2008; Maimon et al., 2010; Schnell et al., 2014), we investigated, whether spontaneous and looming-elicited saccades have a similar effect on HS cells and how they affect the cells’ processing of horizontal motion. In contrast to previous studies, we found a large variability between different HS cells regarding the impact of an efference copy on their activity during both spontaneous and looming-elicited saccades (Fenk et al., 2021; Kim et al., 2017, 2015). If present, the efference copy is able to inhibit responses to preferred direction horizontal motion in HS cells consistent with the idea that it counteracts reafferent sensory input. However, we also found that the looming stimulus itself, when presented on the ipsilateral side, has a similar effect independent of whether the fly performed a saccade or not. Therefore, the processing of optomotor stimuli is not only affected by an efference copy, but also by stimulus history.

## RESULTS

To study the effect of an efference copy on the optomotor pathway, we recorded the membrane potential of HS cells from head-fixed flies during tethered flight, while monitoring turning behavior by measuring the difference between the left and right wing stroke amplitudes (L-R WSA) (Fig. 1) (Maimon et al., 2010; Suver et al., 2016). HS cells in the right optic lobe respond to clockwise motion with a depolarization and to counter-clockwise motion with a hyperpolarization. We recorded from randomly selected HS cells from the right side of the brain only, whose dendritic receptive field therefore cover the right visual hemisphere.

**Figure 1.**
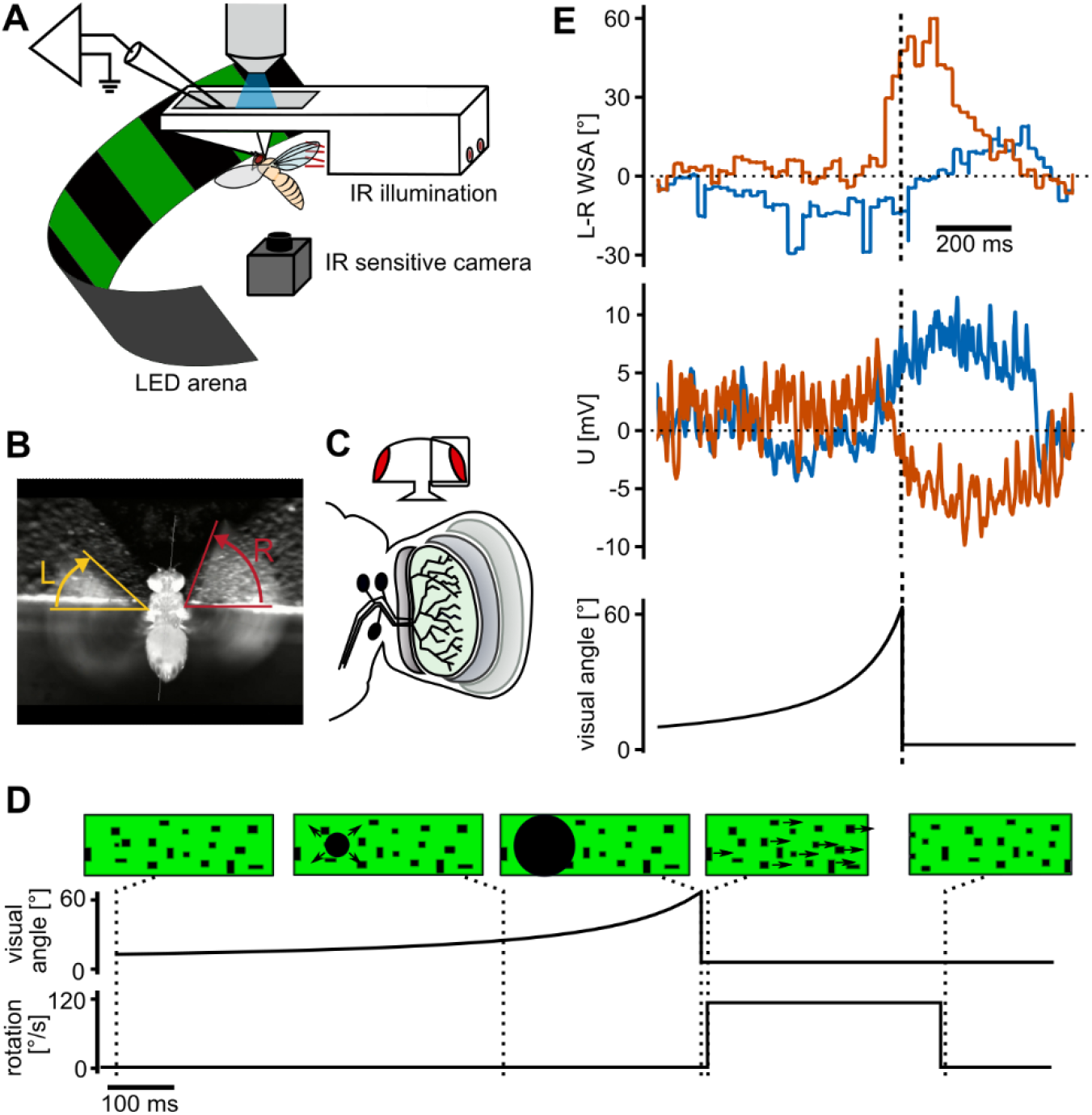
Whole cell recording of HS cells in tethered flying *Drosophila* reacting to looming stimuli. (**A**) Schematic of the setup combining analysis of flight behavior under visual stimulation with whole-cell patch-clamp recordings in head-fixed *Drosophila*. (**B**) Example image of the video stream demonstrating the analysis of wing stroke amplitude. (**C**) Schematic representation of the three HS cells located in the lobula plate within the optic lobe of the fly brain. (**D**) Visual stimulation protocol: a dark spot expands to a disc covering a large part of the visual field to simulate an object approaching at constant speed (looming stimulus) within either the left or right visual hemisphere. In some trials, this stimulus is immediately followed by a rotation of the whole field (called optomotor stimulus). (**E**) Example traces of behavioral response to looming from the left visual field from one fly. In many trials, the fly executed a rapid turn (saccade) away from the stimulated side (red trace, L-R >> 0 corresponds to a rightward turn). The same individual does not perform a saccade in other trials (blue). The membrane potential from the simultaneously recorded HS cells exhibits a hyperpolarization only during trials, where the fly initiates a saccade. Saccades typically occur shortly before the looming disc reaches its maximal size.

### HS cell activity during saccades elicited by ipsilateral looming

To elicit turns, we presented a looming stimulus from either the left (contralateral) or the right (ipsilateral) side using an LED display (Fig. 1D). This stimulus mimicked an object approaching at a velocity of 1.5 m/s and was sufficient to elicit changes in L-R WSA. This change in L-R WSA likely corresponds to a free-flight saccade away from the stimulated side. In the following, we refer to these changes in L-R WSA as saccades, although the flies cannot actually perform a turn under our experimental conditions. Execution of saccades was variable, meaning that flies did not always respond to the looming stimulus with a change of L-R WSA (Fig. 1E). We made use of this variability to test, whether an efference copy of the flies’ behavior influences the membrane potential of HS cells similar to what has been reported for spontaneous saccades (Kim et al., 2017, 2015). We used a saccade detection algorithm on the L-R WSA data from all flies to separate trials with saccades from those without saccades (see Methods and Suppl. Fig. 1) and averaged the HS cell membrane potential and L-R WSA across the two groups.

For looming stimuli presented on the right, which elicits a turn to the left (measured as decrease in L-R WSA) (Fig. 2A), we did not find a qualitative difference in the average HS cell membrane potential between trials with and without a saccade (Fig. 2B). We did notice, however, that after the end of the stimulus presentation, when the looming stimulus disappeared, the membrane potential of the HS cells dropped rapidly and became slightly hyperpolarized independent of whether the flies performed a saccade or not.

**Figure 2.**
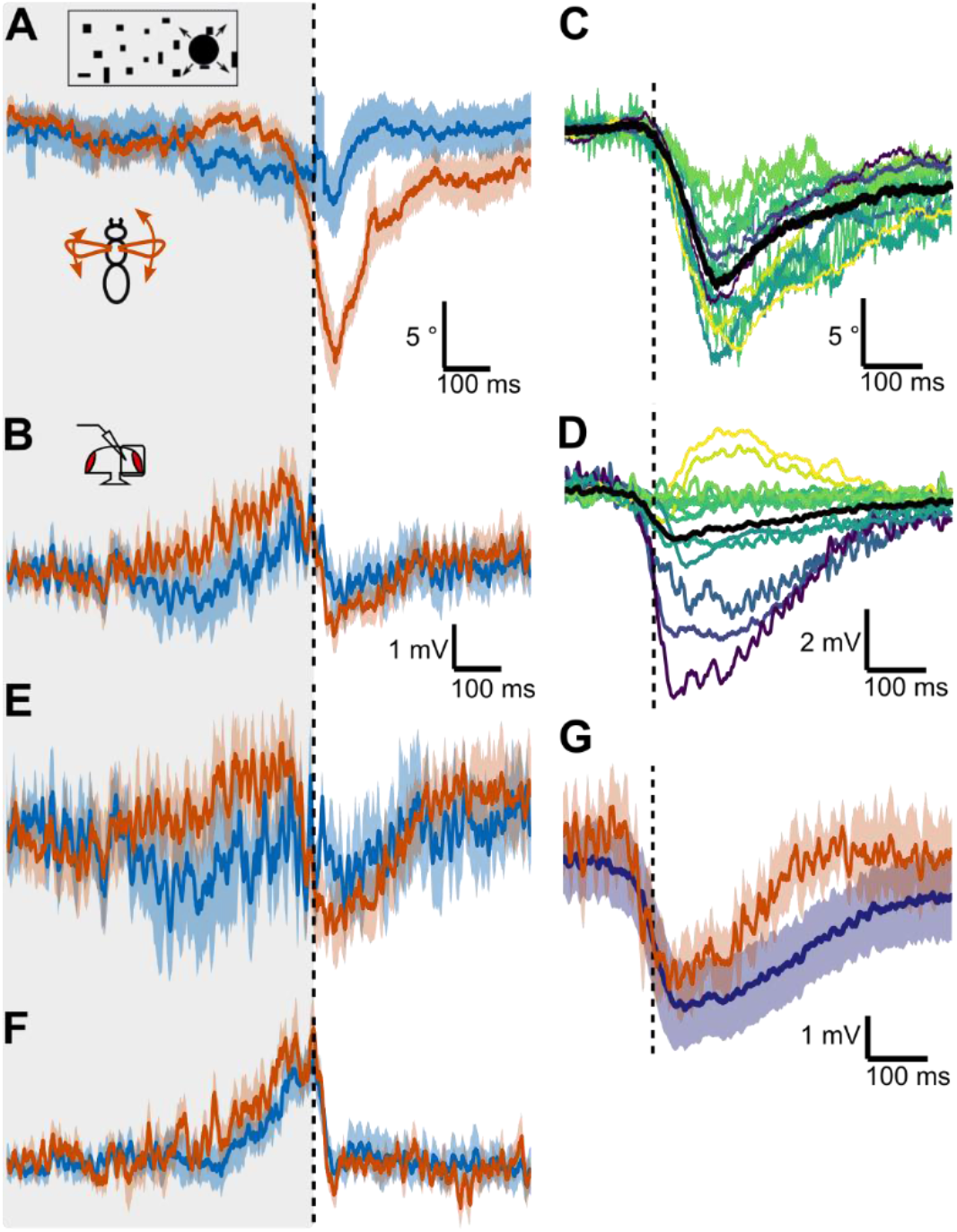
HS cell responses during looming-induced saccades to the left. (**A**) Mean ± s.e.m. of L-R WSA of all flies in trials with looming stimulus presented on the right (ipsilateral) side separated into trials, during which flies reacted with a saccade to the left (red, n=62) or not (blue, n=65), using a wavelet-based algorithm (see Methods and Suppl. Fig. 1). The dotted line marks the end of the looming stimulus. N = 13 flies. (**B**) Simultaneously recorded membrane potential of HS cells averaged across cells as in (A). One cell was recorded per fly. Shown are means ± s.e.m. (**C**) L-R WSA during spontaneous saccades to the left for the same flies as in (A). Each color represents the mean of one fly. The population average in shown in black. (**D**) Mean changes in HS cell membrane potential during spontaneous saccades corresponding to (C). (**E**) Membrane potential of a subset of cells selected based on their hyperpolarizing response during spontaneous saccades (HP_spont_) (see D) from looming trials with an escape saccade (red, n = 25) or without a saccade (blue, n = 21) averaged across flies ± s.e.m (N = 6 cells). (**F**) Average membrane potential ± s.e.m of the remaining cells (NP_spont_ N = 5 cells) as in (E) (n = 18/27 trials). (**G**) Means ± s.e.m. of the cells that hyperpolarized during spontaneous saccades (peak voltage lower than 0, purple, N = 6 cells). For comparison the responses of the same cells during looming-elicited saccades from (C) is replotted in red.

### HS cell activity during spontaneous saccades

It has been reported previously that spontaneous saccades to the left lead to a transient hyperpolarization of HS cells in the right lobula plate (Kim et al., 2015). We detected spontaneous saccades in our recordings using a similar algorithm as for the looming-elicited saccades, but limited to periods, during which visual stimuli were stationary (Fig. 2C). We confirmed that on average HS cells in the right hemisphere hyperpolarize during leftward saccades, although there was a lot of variability across cells with two HS cells even exhibiting a slight depolarization during the peak of the saccade (Fig. 2D). The variability is rooted in neither the detection efficiency nor the amplitude of saccades, given that flies with large saccades figure among the most depolarizing or hyperpolarizing cells.

To test whether spontaneous saccades had a different effect on HS cells than looming-elicited saccades, we sorted our recorded cells into two groups based on whether they exhibited a hyperpolarization during spontaneous saccades (HP_spont_ group, N=6) or no change in polarization (NP_spont_ group, N=5, omitting the depolarizing cells). Cells in the HP_spont_ group also showed a hyperpolarization during looming-elicited saccades (Fig. 2E), which is comparable to spontaneous saccades (Fig. 2G). This is the case despite changes in L-R WSA being larger during looming-elicited saccades. In the NP_spont_ group there was no difference between trials with and without a saccade during the looming stimulus (Fig. 2F). These results indicate that a hyperpolarizing efference copy acts on only a subset of HS cells during both spontaneous and looming-elicited saccades.

Since the HP_spont_ and NP_spont_ subsets also respond differently to the looming stimulus itself (compare Fig. 2E and F), we hypothesized that they may correspond to different HS subtypes. There are three HS cells per hemisphere, which cover different parts of the visual field, and are called HSN (north → dorsal part), HSE (equatorial) and HSS (south → ventral part) (Fig 3A) (Fischbach and Dittrich, 1989; Schnell et al., 2010). However, for the majority of cells shown here, we did not obtain good intracellular dye fills and thus do not know the precise subtype. To test, whether the effect of an efference copy depends on subtype, we drew on data recorded in another context and extracted spontaneous saccades to the left from them (Fig. 3B) (Schnell et al., 2014). Applying the same algorithm to this data with additional selection criteria (see Methods), we obtained saccade-related-potentials in the absence of visual stimulation for 16 HS cells with known subtype. Similar to the case above, saccade-related potentials vary from hyper-to depolarization despite exhibiting strong saccades in a narrow range of amplitudes (Fig. 3C, D). However, results vary between subtypes. Prominently, all HSN cells exhibited a hyperpolarization. In contrast, 3 out of 4 HSS cells exhibited a biphasic response with a weak hyperpolarization at saccade onset, followed by a depolarization that reached its peak just after the peak L-R WSA. HSE cell responses fall in between with some exhibiting a hyperpolarization and some a depolarization. Accordingly, we find the difference in saccade-related-potentials between HSN and HSS to be statistically significant (p < 0.01, 2-sided Mann-Whitney-U test), which is not the case for HSN and HSE (p ≈ 0.08). Altogether, these data confirm the cell-to-cell variability, which can be explained by assuming that an efference copy during saccades has different effects on the different HS cell subtypes.

**Figure 3.**
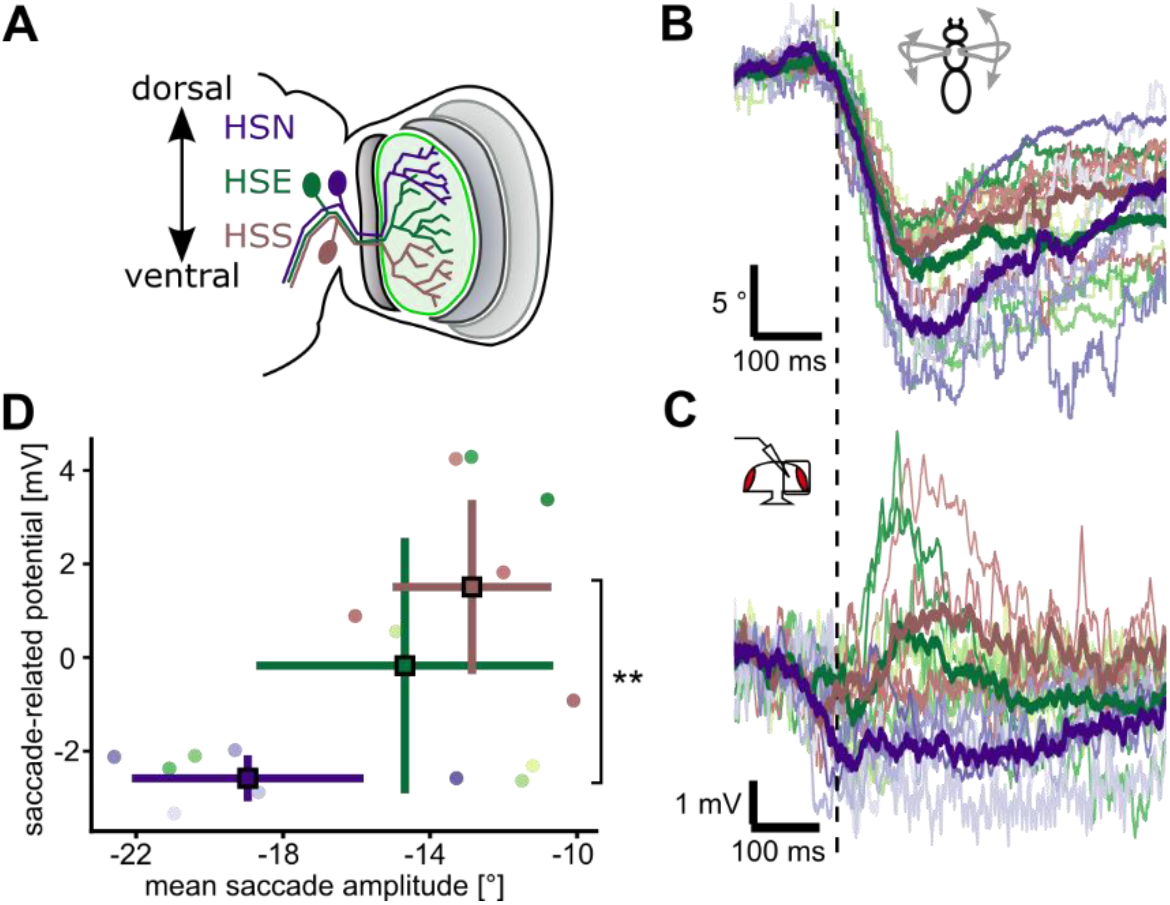
The influence of spontaneous saccades onto different HS cell subtypes. (**A**) Schematic of the three HS subtypes, HSN (purple), HSE (green), HSS (brown). Dendritic receptive fields cover different areas along the dorso-ventral axis with considerable overlap. (**B**) L-R WSA changes classified as spontaneous saccades to the left from previously recorded data. Each line represents the mean for one fly, color-coded according to HS cell subtype as in A. Thick lines represent the mean of the different subtypes. N = 16 (5 HSN, 7 HSE, and 4 HSS cells). (**C**) Mean membrane potential changes of HS cells during spontaneous saccades corresponding to L-R WSA data shown in B. (**D**) Scatter plot of average saccade amplitude and peak voltage change for each cell, color-coded according to subtype. Mean and standard deviation are marked by squares. Membrane potential changes of HSN and HSS are significantly different from each other (p = 0.0095, Mann-Whitney-U 2-sided test), while those of HSN and HSE are not (p = 0.079).

### HS cell activity during saccades elicited by contralateral looming

To test for the effect of saccades in the opposite direction, we presented looming stimuli on the left (contralateral) side, which elicits an increase in L-R WSA corresponding to a rightward saccade in most trials (Fig. 4A), and repeated the same analysis as described for ipsilateral looming. The membrane potential of the recorded HS cells became on average slightly hyperpolarized during trials with a saccade compared to trials without (Fig. 4B). This hyperpolarization occurred after the presentation of the stimulus, when the L-R WSA reached its peak. We also see the effect, when comparing trials from just one fly, suggesting that this is not simply due to variability between different flies (Fig. 1E). Thus, there is evidence that an efference copy influences the contralateral HS cell membrane potential during visually elicited saccades.

**Figure 4.**
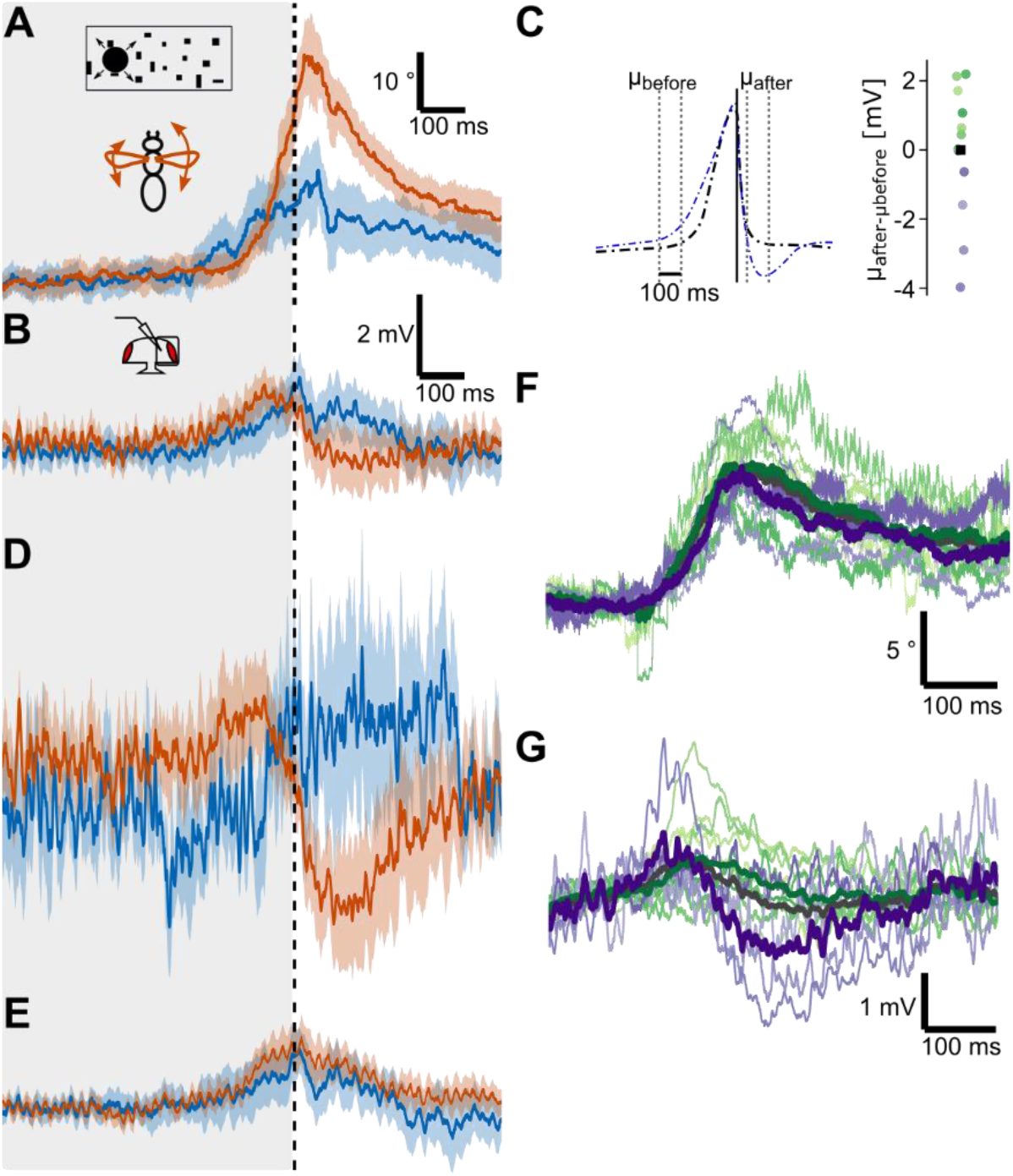
HS cell responses during looming-induced saccades to the right. (**A**) Mean L-R WSA ± s.d. in response to looming on the contralateral side (with respect to recorded cell) averaged across trials with a detected saccade (red, n = 74 trials) and those without (blue, n = 53 trials). N = 13 flies. (**B**) HS cell membrane potential changes corresponding to the L-R WSA data shown in A. One cell was recorded from each fly. (**C**) Criterion to split HS cells into groups according to their response to the looming stimulus during tethered flight. The mean membrane potential after the stimulus is subtracted from a baseline in the middle of the expansion (300 ms before maximal size). Flies are grouped by whether this difference is below (purple dots, HP_contra_) or above average (green dots, NP_contra_). (**D**) Membrane potential of the HP_contra_ subset of HS cells (purple cells from C, N = 4) for trials with (red, n = 23) and without saccades (blue, n = 8). Shown here is the mean ± s.d. across all trials from all flies because of too few non-saccade trials per fly. (**E**) Same as (D), but for NP_contra_ cells (green dots from C, N = 9 flies, n=51/45 trials). (**F**) L-R WSA during spontaneous saccades to the right for the same flies as in (A). Each color represents the mean of one fly. Thick lines represent averages across the different subsets according to (C) and across all flies (grey). (**G**) Membrane potential of HS cells during spontaneous saccades corresponding to (F).

When looking at individual flies, we did not find the hyperpolarization during rightward saccades in all of the cells we recorded from. Rather, we were able to separate our recordings into two groups based on whether we were able to observe the effect (HP_contra_) or not (NP_contra_) (Fig. 4C). In four HS cells from four different flies we observed a clear hyperpolarization (Fig. 4D). In the remaining nine cells, we saw no clear difference in HS cell membrane potential between trials with and without a saccade (Fig. 4E).

The hyperpolarization we observed is different from the effect reported during spontaneous saccades. For spontaneous saccades, a depolarization has been reported during rightward turns (Kim et al., 2015). We also observed a slightly stronger depolarization during the presentation of the looming stimulus in trials with a saccade compared to those without (Fig. 4D), which occurred before L-R WSA started to rise. Because the depolarization could also be caused by the visual stimulus via input from contralateral lobula plate tangential cells for example, we cannot conclusively say, whether it is influenced by an efference copy or not. When we averaged the HS cell membrane potential during spontaneous saccades to the right of only the HP_contra_ group, we observed a similar hyperpolarization (Fig. 4F, G). This hyperpolarization is again preceded by a small depolarization suggesting that both are indeed the consequence of an efference copy. Note that the cells contained in the HP_contra_ group also hyperpolarize during saccades elicited by ipsilateral looming (HP_spon_, see Table 1) and therefore likely represent HSN and/or HSE cells as well. Altogether, there is no qualitative difference between the effect of spontaneous and looming-elicited saccades onto HS cells, although there is substantial cell-to-cell variability. In conclusion, for looming stimuli presented on the contralateral side, we did find evidence for a hyperpolarizing efference copy acting on a subset of HS cells.

**Table 1.** Summary of classification of recorded cells according to their change in membrane potential during spontaneous saccades away from the ipsilateral side (HP_spont_/NP_spont_, see Fig. 2) and according to their response to a contralateral looming stimulus (HPcontra/NPcontra, see Fig. 4). Groups are highly overlapping, with HP_contra_ being a subset of HP_spont_ and NP_spont_ a subset of NP_contra_. HP = hyperpolarized, NP = no change in polarization.

|  |  | Contralateral looming |  |
| --- | --- | --- | --- |
|  |  | HP<sub>contra</sub> (N = 4) | NP<sub>contra</sub> (N = 9) |
| Ipsilateral<br>looming | HP<sub>spont</sub> (N = 6) | 4 | 2 |
|  | NP<sub>spont</sub> (N = 5) | 0 | 5 |
|  | Depolarizing (N = 2) | 0 | 2 |

### Impact of looming-elicited saccades onto processing of horizontal motion stimuli

A major advantage of our approach, compared to studying spontaneous saccades, is that we can elicit the behavior and thus study its effect on a subsequent optomotor stimulus. To study, whether motion responses are altered in HS cells after a saccade, we presented a large-field horizontal motion stimulus directly after the looming stimulus, when the peak of the saccade occurred (Fig. 1D). Note that we chose stimulus parameters that are slower than what flies would perceive during a free flight saccade to drive strong HS cell responses. This stimulus pattern consisted of random dots moving either clockwise or counter-clockwise. As a comparison, we measured responses to just the moving dots without the preceding looming stimulus. A looming stimulus presented on the right triggers a saccade to the left, which during free flight would lead to clockwise motion, which depolarizes the HS cells in the right optic lobe. We observed that the response to clockwise motion after the looming stimulus was rising much slower when compared to the response to the motion stimulus presented alone (Fig. 5A, B). However, this was the case during flight saccades (red) as well as during non-flight trials (green). Further, the cells in the NP_spont_ group, which did not show any hyperpolarization to the looming stimulus (Fig. 2E), have a similarly suppressed motion response (Fig. 5B).

**Figure 5.**
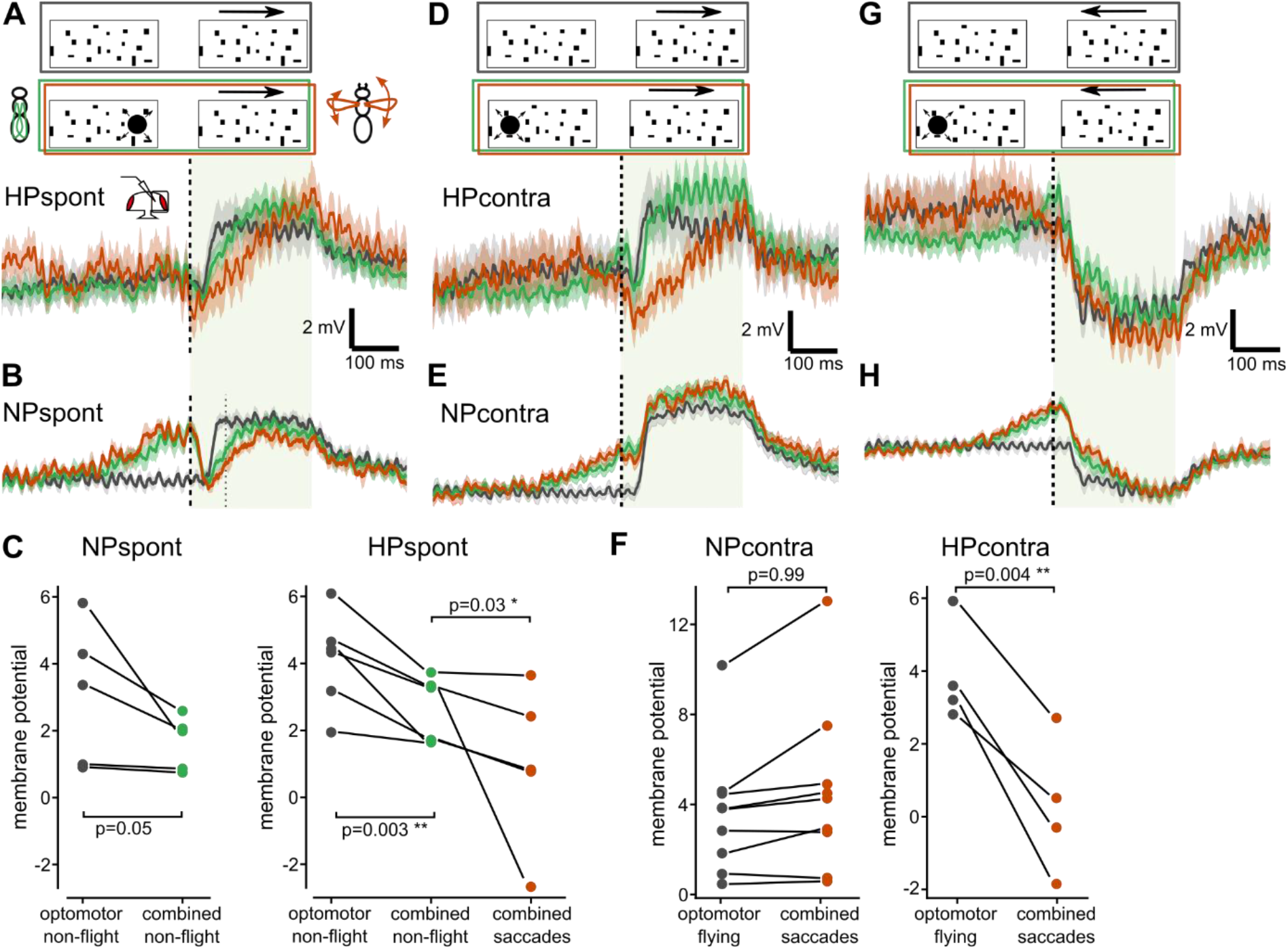
Processing of horizontal motion after a looming stimulus in HS cells. All traces represent mean ± s.e.m. membrane potential. (**A**) Response to an excitatory (clockwise) motion stimulus after a looming stimulus on the ipsilateral side of cells that hyperpolarize during saccades (HP_spont_, same subset as in Fig. 2E, N = 6). Trials with saccades are shown in red (n = 19), non-flight trials in green (n = 70) and responses to the optomotor stimulus without preceding looming in grey (n = 50). (**B**) Same as (A), but for NP_spont_ (subset of HS cells that do not hyperpolarize, see Fig. 2D, N = 5, n = 24/62/48). (**C**) Average change in membrane potentials during the early phase of the optomotor stimulus (80 ms after looming, grey dotted line in B) for each cell in the NP_spont_ and HP_spont_ group respectively. Color code as in A. A paired t-test was used for comparing responses to the combined stimulus during non-flight with the optomotor stimulus, a Wilcoxon paired sign test for comparing trials with saccades during the combined stimulus with the opotmotor stimulus only. (**D**) Response to an excitatory (clockwise) motion stimulus after a looming stimulus presented in the contralateral (left) visual hemisphere of cells that hyperpolarize during saccades HP_contra_ (same subset as in Fig. 4D, N = 4, n = 24/45/28). Color code as in (A). (**E**) Same as (D), but for NP_contra_ (subset of HS cells that do not show a hyperpolarization, see Fig. 4E, N = 9, n = 41/115/101). (**F**) Average membrane potential for each cell as in (C). A Wilcoxon paired sign test was used for the NP_contra_ cells and a paired t-test for the HP_contra_ cells (Shapiro-Wilk normality test, p=0.99)) (**G**) Response to an inhibitory (counter-clockwise) motion stimulus after a looming stimulus on the contralateral side of HP_contra_ (same subset as in panel D, n = 25/43/29). Color code as in (A). (**H**) Same as (G), but for NP_contra_ subset (same subset as in panel E, n = 48/109/105).

We quantified this response by averaging the membrane potential during the early phase of the optomotor stimulus (Fig. 5C). Considering all cells together, the response during the combined looming and optomotor stimulus during non-flight was reduced very significantly compared to the presentation of the optomotor stimulus alone (p = 1.6*10^-3^, 2-sided t-test; p-values of individual groups marked in Fig. 5C). We therefore conclude that this suppression of the response to horizontal motion is induced by the looming stimulus itself presented within the dendritic receptive field of the recorded HS cells and not by the flies’ behavior. Therefore, in the case of an ipsilateral looming stimulus, an efference copy might not be necessary to suppress reafferent responses to panoramic motion that could interfere with the execution of the saccade. However when the behavior is executed, in HP_spont_ cells (hyperpolarizing during saccades towards the left) the response to clockwise (rightward) motion is even further suppressed compared to non-flight trials, showing that the presumed efference copy has an additional effect on motion processing (Fig. 5A and C) (p = 0.031, 1-sided Wilcoxon paired sign test).

For the looming stimulus presented on the left or contralateral side, we again plotted averaged responses separately for HP_contra_ cells (which showed the hyperpolarization) and NP_contra_ cells (which did not) (Fig. 5D, E). Here we found that responses to clockwise motion were again rising significantly slower after a saccade compared to the response to just the wide-field motion stimulus, but only in the HPcontra cells where we found evidence for the hyperpolarizing efference copy, and only during saccades (p = 0.0042, 1-sided paired t-test) (Fig. 5F). Responses to counter-clockwise motion, which hyperpolarizes the recorded HS cells, were not strongly affected during saccades (Fig. 5G, H). This finding suggests that the efference copy acting on HS cells during a contralateral looming stimulus serves to suppress responses to motion that would normally depolarize the cell.

## DISCUSSION

In the optomotor pathway of *Drosophila*, an efference copy has been shown to act on HS cells during spontaneous saccades (Kim et al., 2017, 2015). HS cells compute information about large-field horizontal motion elicited by self-rotation of the fly and are thought to mediate compensatory optomotor responses. Therefore, their activity could interfere with the self-elicited turn.

Here, we show that looming stimuli eliciting evasive saccades have a similar effect on HS cells as spontaneous saccades (Figs. 2 and 4). In contrast to the previous studies, however, we found a large cell-to-cell variability within the recorded HS cells with respect to the action of an efference copy. In addition, we observed that excitatory responses to preferred-direction horizontal motion in HS cells are suppressed after looming stimuli on the ipsilateral side of the recorded cell even in non-flying flies and thus independent of the execution of a saccade (Fig. 5). This suppression is therefore likely a consequence of the looming stimulus itself and not of an efference copy of the motor command controlling the saccade, although the latter appears to play an additional role in a subset of HS cells. We can only speculate, whether this is a specific effect of the looming stimulus through a feedback mechanism or a consequence of light or motion adaptation in response to the dark disc.

After a looming stimulus on the contralateral side, we also observed a hyperpolarizing response only in a subset of HS cells, which was dependent on the execution of a saccade and thus likely mediated through an efference copy (Fig. 4). This efference copy was able to suppress responses to a subsequent excitatory motion stimulus, but did not affect an inhibitory motion stimulus (Fig. 5). This finding is surprising given that a saccade towards the contralateral side is expected to elicit null-direction horizontal motion and therefore to hyperpolarize HS cells. While a hyperpolarization of HS cells has been shown to be able to elicit behavioral responses during walking (Busch et al., 2018), our findings suggest that predominantly excitatory responses of HS cells are suppressed during flight saccades. However, we did not study motion responses during the early phase of the saccade, when the looming stimulus was still present, so we cannot exclude that inhibitory responses are affected during that time.

We also found that the cell-to-cell variability with respect to the action of an efference copy can partially be explained by taking the different subtypes of HS cells into account (Fig. 3). The hyperpolarization during spontaneous saccades towards the contralateral side was predominantly observed in HSN cells and some HSE cells, while HSS cells even showed a depolarization. A recent study found evidence for an efference copy acting on HS cells during saccades elicited by looming stimuli presented on the ipsilateral side of the recorded cells (Fenk et al., 2021). Fenk et al. did not report a similar variability between cells as our study, which is likely due to the fact that they only recorded from HSN cells. Although our data on looming-elicited saccades did not discriminate between HS subtypes, the correlation between responses to spontaneous and looming-elicited saccades suggests that predominantly HSN and some HSE cells receive an hyperpolarizing input during escape saccades. While we generally confirm the results of their study, we find that the effect of both the stimulus as well as saccades onto HS cells is surprisingly complex, which has implications for the search of the neurons that mediate the efference copy.

What might explain the large cell-to-cell variability we observed regarding the effect of saccades onto HS cells? Because of the complexity of these flight maneuvers, we do not know how the different types of HS cells would respond to the visual motion stimulus that would be caused by the fly’s self-motion during a saccade in free flight. It is possible that the different types of HS cells because of their different receptive fields (Krapp et al., 2001; Schnell et al., 2010) could be differently affected by the motion stimulus during a saccade. Therefore, an efference copy might only act on some cells and not others. Alternatively, different HS cell subtypes might have different functions in controlling flight behavior and therefore could be differently affected by an efference copy during saccades. In contrast to HSN and HSE cells, which receive input from heterolateral LPTCs, for example, HSS is not responding as much to back-to-front motion on the contralateral side (Hausen, 1982a; Schnell et al., 2010) and might therefore play a role in regulating forward motion rather than turning. Our data are consistent with a circuit, in which HSN receives inhibitory and HSS excitatory input during saccades towards the contralateral side. Because of the electrical coupling between HS cells (Haag and Borst, 2005, 2002; Schnell et al., 2010), HSE could receive indirect input via HSN and HSS with one or the other dominating in some cells, which might explain the response variability we observe in our recordings. Alternatively, there may be subtle differences in the behavior that we are not aware of, as we only measure L-R WSA. These differences in behavior could lead to different effects onto HS cells. Further research is needed to discriminate between these different possibilities.

A saccade during free flight is a complex maneuver consisting of a roll, a counter-roll and a yaw turn to realign the body with the new heading (Muijres et al., 2015, 2014; Schilstra and Hateren, 1999). It is therefore difficult to judge, what the changes in L-R WSA correspond to. During free flight saccades, changes in L-R WSA were shown to be small and seem to have more impact on roll than on yaw torque, suggesting that the changes in L-R WSA we measure during tethered flight could correspond to the roll maneuver (Dickinson and Muijres, 2016). It has been shown that during looming-elicited escape saccades in free flight the fly sacrifices stability for speed (Muijres et al., 2015, 2014). Spontaneous saccades on the other hand are more stereotyped and the yaw maneuver happens with smaller delay, such that times of motion blur are minimized (Land, 1999). The quantitative difference we find between spontaneous and looming-elicited saccades might therefore be due to difference in the motor programs underlying these two behaviors. While a descending neuron has been described that is active during changes in L-R WSA associated with spontaneous as well as looming-elicited saccades (Schnell et al., 2017), it is still unclear, which neural substrate could mediate the efference copy to the HS cells.

Especially for flying insects, vision is an essential modality, to which a sizeable portion of their brain is dedicated. The processing of visual information, however, needs to depend on the context, such as the behavioral and motivational state of an animal or the presence of other stimuli, to lead to the appropriate behavioral response. Looming stimuli for example can signal an approaching predator or an impending collision. Illustrating the dependence on behavioral context, they can elicit escape jumps in stationary flies, whereas in a flying fly they can trigger either a landing response or an evasive turn (Ache et al., 2019; Bender and Dickinson, 2006; Tammero and Dickinson, 2002; Tammero and Dickinson, 2002; von Reyn et al., 2014).

Efference copies and corollary discharge signals have been shown to influence sensory processing in a variety of different systems and at different levels culminating in the predictive coding hypothesis (Combes et al., 2008; Crapse and Sommer, 2008; França de Barros et al., 2022; Fukutomi and Carlson, 2020; Poulet and Hedwig, 2006; Sommer and Wurtz, 2008; Wurtz, 2008). Despite decades of research, the mechanisms underlying the action of efference copies remain largely speculative. In *Drosophila*, the genetic tools as well as the recording techniques will give as the chance to dissect the circuitry underlying efference copies, which could provide fundamental insights into the way nervous systems work.

Here, we show that in *Drosophila* an efference copy is able to suppress responses to panoramic visual motion in the optomotor system during self-elicited turns, however, with a large cell-to-cell variability. Our study supports the notion that information processing already at early stages, i.e. at the level of sensory systems, is strongly influenced by stimulus history as well as behavioral state. The more important it is to consider the behavioral context when studying nervous system function.

## MATERIALS AND METHODS

### Resource availability

Raw data are available under DOI:10.17617/3.92. Further information and requests for resources and reagents should be directed to and will be fulfilled by the lead contact, Bettina Schnell.

### Code

The source code used for analysis will be made available as GitHub repository and a unique DOI will be provided by the time of publication.

### Fly stocks

Strains of *Drosophila melanogaster* were bred on premixed cornmeal-agar medium (JazzMix Drosophila food, Fisherbrand) at 25°C on a 12h day/12h night cycle. Flies used for experiments were of either wild-type CantonS (two flies) or expressing GFP in HS cells using the Gal4/UAS system (GMR81G07-GAL4 or GMR27B03-GAL4). All individuals had at least one wild-type allele for the *white* gene. To avoid side-effects of in-breeding, GMR27B03-GAL4; 2xUAS-EGFP flies were backcrossed to wildtype flies (Canton S and Top Banana), and offspring selected for fluorescence every few generations. We used female flies aged 2 to 5 days after eclosion for all experiments.

### Electrophysiology

*In vivo* patch-clamp recordings of HS cells were performed in whole-cell configuration in current-clamp mode. Flies were anaesthetized on a cold-plate, legs amputated, and fixed by the head to custom-made pyramidal holders (Maimon et al., 2010) that allow access to portions of the head while offering free space to the wings using UV-curing glue. The head was bent forward to expose the posterior part in the surgery window of the holder. The head capsule was covered with extracellular saline (prepared according to Wilson and Laurent (Wilson and Laurent, 2005), in mM: 103 NaCl, 4 MgCl_2_, 3 KCl, 1.5 CaCl_2_, 26 NaHCO_3_, 1 NaH_2_PO_4_, 10 Trehalose, 10 Glucose, 2 Sucrose and 5 TES, adjusted to 275 mOsm) and cut open using a syringe needle. The brain was perfused with extracellular saline bubbled with 95% O_2_/5% CO_2_ to reach pH 7.3. The neurolemma and extracellular matrix covering the lobula plate was digested by applying Collagenase (0.5mg/mL Collagenase IV in extracellular saline, Worthington) through a low-resistance pipette (BF150-110-10, Sutter Instruments). HS cells were targeted under visual control with a 40x immersion objective (Nikon NIR Apo 40x/0.80W), phase-contrast filter, IR lighting (850 nm, Thorlabs) and CMOS camera (Thorcam Quantalux, Thorlabs). GFP-expression was visualized by a brief excitation of eGFP in transgenic flies. Patch pipettes were pulled with a P-1000 Flaming/Brown puller (Sutter Instruments) from thick-walled capillaries (BF150-86-10, Sutter Instruments), adjusted to a resistance of 6 to 8 MΩ. Intracellular solution was prepared according to (Wilson and Laurent, 2005) (in mM: 140 mM potassium aspartate, 1 mM KCl, 10 mM HEPES, 1 mM EGTA, 4 mM MgATP, 0.5 mM NaGTP, 0.5% biocytin, adjusted to pH 7.3 and 265 mOsm) plus AlexaFluor-568. Signals were recorded with a BA-01X bridge amplifier (npi electronics) and a BNC-2090A digitizer (National Instruments) with MATLAB (version R2017b, The MathWorks). Recordings from HS cells presented in Fig. 3 were performed using the flies and setup described previously (Schnell et al., 2014).

### Behavioral monitoring

The flight behavior was observed by an IR-sensitive camera (Basler acA645-100gm equipped with a 1.0X 94mm WD InfiniStix lens, Infinity Photo-Optical) placed below the fly and illuminating the wings from the fly’s posterior with the IR LEDs mentioned above. The front edge of the wing was detected from the image stream using the Kinefly software (Suver et al., 2016) running on ROS (version Kinetic Kame), which outputs a signal that was fed into the same digitizer as the electrophysiological signal. An important change has been made to the Kinefly wing detection algorithm. The front edge of the wing was determined as the first threshold crossing of a derivative luminance signal extracted from the image, which made the detection less error-prone.

### LED arena and visual stimulation

The fly was at the center of a cylindrical arena of LED panels (IO Rodeo) as described (Reiser and Dickinson, 2008). Our arena is made of monochromatic LEDs (565 nm) where each pixel subtends an angle of 2.25 ° on the fly eye, covering a total visual field of about 195° in azimuth and 60° in elevation. Here, we used a frame shift rate of 50 Hz, while the arena was tuned to a flicker rate of about 744 Hz. The visual scene presented to the fly consisted of dark squares on a bright background with maximal contrast. Centered at an angle of 54° on either the left or right side, a dark disc expanded, simulating an object approaching at constant speed of 1.5 m/s. Thus the visual angle covered by the imaginary object is given by 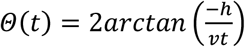 with h = 0.1 m, v = 1.5 m/s and t = 0 at the expected moment of impact (Gabbiani et al., 1999). The maximal lateral extent of the stimulus was 76.5°. At time t’ > 0, the looming disc disappears. The background consisted of randomly distributed dark rectangles on a bright background with 100% nominal contrast, with each rectangle measuring at least 5° x 5° in the visual field. This background was rotated with a speed of 112.5 °/s for 240 ms either clockwise or counter-clockwise, which we call optomotor stimulus. In a given trial, the stimulus consisted of either looming (left/right) only, background rotation (either direction) only, or looming followed by background rotation, making for a total of eight different stimulus combinations. Each trial lasted for 1.35 s, with an inter-trial interval of 3 s. The order of these stimuli was permuted. Each fly completed several experimental runs.

### Quality criteria

All data analysis was performed in Python (version 3.7). To make sure that we would be able to detect a hyperpolarizing effect of an efference copy, we excluded HS cell recordings, if they did not respond to a square-wave pattern rotating counter-clockwise by a tonic hyperpolarization < −2 mV. All experimental runs for one fly were gathered and divided into trials, including 100 ms before and 250 ms after the actual trial. An angular threshold was applied to the wing stroke amplitude data of both wings to avoid contamination of the data by misdetections due to reflection artifacts. This angle depended on preparation and light conditions, but in most instances was < 8°, which does not occur in normal flight. If both wings were consistently below threshold, the trial was classified as non-flying. If there was a discrepancy between left and right wing (above/below threshold) for more than 30 ms in a trial, corresponding to less than two detection frames, the trial was discarded. Spontaneous saccades were extracted from data during the inter-trial intervals, where the visual input consisted of the static background only. In the older datasets (recorded as described in (Schnell et al., 2014)), the experimental data were truncated where resting membrane potential started to drift more than ≈ 0.01 mV/s.

### Saccade detection

Saccades were detected using an algorithm based on continuous wavelet transform (CWT) applied to a differentiated signal without relying on low- or high-pass filtering to avoid boundary artifacts in shorter flight bouts. Briefly, any signal can be decomposed into a linear sum of many smaller wavelet functions, which have a limited width in the time and frequency domains. An abrupt event of significant amplitude shows in CWT as a time-localized peak over a broad range of wavelet scales. Thus, it is detectable by tracking the uninterrupted peaks in the CW-transformed signal over scales. We use the algorithm developed by Du et al. (Du et al., 2006), as implemented in the scipy.signal (scipy version 1.6.2) package. First, to simplify saccade detection, we downsampled the noise-corrupted piece-wise-constant data obtained by recording the wing stroke amplitude together with electrophysiological data through a fitting procedure to obtain a single value per analysis frame. The downsampled signal was median-filtered with filter width 3 to remove single outliers, and then differentiated using the central difference formula with a half-width of 50 ms, to build the detection signal. These data were fed into the peak-detection function with scales {2^2+j/16^}, j = {0,2,…,15}, window size of 10 frames, gap threshold of 1, and noise percentage of 5, which constitute numerical values. To detect saccades to the left, the negative of the signal was used. Spontaneous saccades, where the direction of the saccade is not determined by the stimulus, were detected in the same way on both the positive and negative signal with a window size of 20 frames and noise percentage of 10 to be more restrictive. For each putative saccade we calculated the peak and onset time and peak amplitude. The peak time was determined by moving forward from the putative peak and finding the first occurrence of crossing 0 (indicating a local extremum) in the signal used for detection and adding the half-width of the differentiation. The onset time was defined as the first minimum before the putative peak in the simple differential (not central difference) of the downsampled original signal. Peak amplitude was calculated by taking the average L-R WSA in the interval of peak time +/− 30ms. To eliminate cases, where the falling slope of a previous saccade was detected as a saccade in the opposite direction, we removed candidate saccades if another deviation from baseline was detected within 160 ms preceding the onset, which exceeded the following percentage of the peak amplitude. For spontaneous saccades, that percentage was 60%, while we chose 75% for saccades during looming trials. Trials were then classified according to whether they contained a saccade that was synchronized to the time when the looming stimulus reached its maximum expansion within a tolerance window of [-200 ms; 60 ms]. As a consequence of these steps, trials classified as non-saccades mostly consist of slow deviations to either side, consecutive turns to unclear direction, and very rarely contained very early saccades (long before the looming stimulus ends).

Additional criteria for the selection of saccades were applied to the data for different HS cell subtypes, shown in Fig. 3, to further reduce the rate of false positives. First, a threshold was applied to the saccade amplitudes, tuned to 4.7, 7.4, 8.8 or 11.8°, where the two highest values were by far the most common. Further, putative saccades were eliminated if there were less than 500 ms between the onset of saccades in opposite directions or if a deviation from baseline before putative saccade onset exceeded 65% of saccade amplitude. Finally a candidate saccade was accepted only if the onset preceded the peak in the derivative signal by at most 100 ms and the peak of the saccade followed by at most 300 ms. This procedure removed a majority of candidates, and the resulting putative saccades were visually verified.

### Experimental design and statistical analyses

No statistical tests were used to predetermine sample sizes, which were, however, in the range of other publications in the field. Detailed numbers including ‘N’ of individual animals and ‘n’ of trials are given in every figure legend. Electrophysiology data of each trial were baseline-corrected to the mean potential in the window of 400 ms from the beginning of the trial. This window contains no responses to the looming stimulus. The data from trials were in most cases averaged per fly, before calculating an average of averages +/− s.e.m. In cases of low number of trials per fly in certain subsets, all trials were instead treated as drawn from the same distribution, and therefore the mean ± s.d. was given. Which of these methods was applied, is stated in the figure captions.

For a statistical analysis, we quantified the response to the panoramic motion stimulus under the different conditions (see Fig. 5) by averaging the baseline-corrected membrane potential for each fly in a window of 80 ms +/− 8 ms after the looming stimulus reached its maximal extension. A Shapiro-Wilk-test was performed to ascertain normality of the distribution with significance threshold p = 0.10, followed by a dependent t-test for paired samples. If the distribution was non-normal, a Wilcoxon’s paired sign test was applied. Clarifications on which test was used is given in figure captions.

Statistical significance of the results on subtype-specificity of responses during spontaneous saccades (see Fig 3D) was determined by 2-sided Mann-Whitney-U tests.

### Criteria for subgroups

First, the difference in membrane potential before and after the looming stimulus was calculated on the mean membrane potential U for each fly for trials with looming stimulus only (no background rotation).

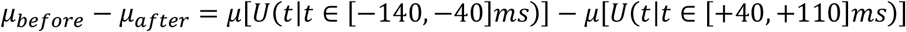

Flies were then grouped in the hyperpolarizing category if that difference was lower (more negative) than the average difference.

## AUTHOR CONTRIBUTIONS

Conceptualization: B.S.; Methodology: P.F., B.S.; Software: P.F.; Formal Analysis: P.F.; Investigation: P.F., B.S.; Writing (Original Draft): P.F., B.S.; Writing (Review & Editing): P.F., B.S.; Visualization: P.F.; Funding Acquisition: B.S.; Supervision: B.S.; Project Administration: B.S.

## ACKNOWLEDGMENTS

We want to thank Tim Krause for maintaining fly stocks, Gaia Tavosanis, Eugenia Chiappe and Michael Dickinson for sharing fly stocks and Elhanan Ben-Yishay and Kevin Briggman for comments on the manuscript. This work was funded by the German Research Foundation (DFG) through the Emmy-Noether program.

## REFERENCES

Ache, J.M., Namiki, S., Lee, A., Branson, K., Card, G.M., 2019. State-dependent decoupling of sensory and motor circuits underlies behavioral flexibility in Drosophila. Nat. Neurosci. 22, 1132–1139.

Bender, J.A., Dickinson, M.H., 2006. Visual stimulation of saccades in magnetically tethered Drosophila. J. Exp. Biol. 209, 3170–3182.

Boeddeker, N., Lindemann, J.P., Egelhaaf, M., Zeil, J., 2005. Responses of blowfly motion-sensitive neurons to reconstructed optic flow along outdoor flight paths. J. Comp. Physiol. A 191, 1143–1155.

Busch, C., Borst, A., Mauss, A.S., 2018. Bi-directional Control of Walking Behavior by Horizontal Optic Flow Sensors. Curr. Biol. 28, 4037–4045.e5.

Combes, D., Le Ray, D., Lambert, F.M., Simmers, J., Straka, H., 2008. An intrinsic feed-forward mechanism for vertebrate gaze stabilization. Curr. Biol. 18, R241–R243.

Crapse, T.B., Sommer, M.A., 2008. Corollary discharge across the animal kingdom. Nat. Rev. Neurosci. 9, 587–600.

Cullen, K.E., 2004. Sensory signals during active versus passive movement. Curr. Opin. Neurobiol. 14, 698–706.

Dickinson, M.H., Muijres, F.T., 2016. The aerodynamics and control of free flight manoeuvres in Drosophila. Philos. Trans. R. Soc. B Biol. Sci. 371, 20150388.

Du, P., Kibbe, W.A., Lin, S.M., 2006. Improved peak detection in mass spectrum by incorporating continuous wavelet transform-based pattern matching. Bioinformatics 22, 2059–2065.

Fenk, L.M., Kim, A.J., Maimon, G., 2021. Suppression of motion vision during course-changing, but not course-stabilizing, navigational turns. Curr. Biol. 31, 4608–4619.e3.

Fischbach, K.F., Dittrich, A.P.M., 1989. The Optic Lobe of Drosophila-Melanogaster. 1. A Golgi Analysis of Wild-Type Structure. Cell Tissue Res. 258, 441–475.

França de Barros, F., Bacqué-Cazenave, J., Taillebuis, C., Courtand, G., Manuel, M., Bras, H., Tagliabue, M., Combes, D., Lambert, F.M., Beraneck, M., 2022. Conservation of locomotion-induced oculomotor activity through evolution in mammals. Curr. Biol. 32, 453–461.e4.

Fujiwara, T., Cruz, T.L., Bohnslav, J.P., Chiappe, M.E., 2017. A faithful internal representation of walking movements in the Drosophila visual system. Nat. Neurosci. 20, 72–81.

Fukutomi, M., Carlson, B.A., 2020. A History of Corollary Discharge: Contributions of Mormyrid Weakly Electric Fish. Front. Integr. Neurosci. 14. 10.3389/fnint.2020.00042

Gabbiani, F., Krapp, H.G., Laurent, G., 1999. Computation of Object Approach by a Wide-Field, Motion-Sensitive Neuron. J. Neurosci. 19, 1122–1141.

Götz, K.G., 1968. Flight control in Drosophila by visual perception of motion. Kybernetik 4, 199–208.

Haag, J., Borst, A., 2005. Dye-coupling visualizes networks of large-field motion-sensitive neurons in the fly. J. Comp. Physiol. A. 191, 445–454.

Haag, J., Borst, A., 2002. Dendro-dendritic interactions between motion-sensitive large-field neurons in the fly. J. Neurosci. 22, 3227–3233.

Haag, J., Borst, A., 2001. Recurrent Network Interactions Underlying Flow-Field Selectivity of Visual Interneurons. J. Neurosci. 21, 5685–5692.

Haikala, V., Joesch, M., Borst, A., Mauss, A.S., 2013. Optogenetic Control of Fly Optomotor Responses. J. Neurosci. 33, 13927–13934.

Hausen, K., 1982a. Motion sensitive interneurons in the optomotor system of the fly. 1. The Horizontal Cells - structure and signals. Biol. Cybern. 45, 143–156.

Hausen, K., 1982b. Motion sensitive interneurons in the optomotor system of the fly. 2. The Horizontal Cells - receptive-field organization and response characteristics. Biol. Cybern. 46, 67–79.

Heisenberg, M., Wolf, R., 1979. On the fine structure of yaw torque in visual flight orientation of Drosophila melanogaster. J. Comp. Physiol. A 130, 113–130.

Joesch, M., Plett, J., Borst, A., Reiff, D.F., 2008. Response properties of motion-sensitive visual interneurons in the lobula plate of Drosophila melanogaster. Curr.Biol. 18, 368–374.

Kim, A.J., Fenk, L.M., Lyu, C., Maimon, G., 2017. Quantitative Predictions Orchestrate Visual Signaling in Drosophila. Cell 168, 280–294.e12.

Kim, A.J., Fitzgerald, J.K., Maimon, G., 2015. Cellular evidence for efference copy in Drosophila visuomotor processing. Nat. Neurosci. 18.

Krapp, H.G., Hengstenberg, R., Egelhaaf, M., 2001. Binocular contributions to optic flow processing in the fly visual system. J. Neurophysiol. 85, 724–734.

Land, M.F., 1999. Motion and vision: why animals move their eyes. J. Comp. Physiol. A 185, 341–352.

Leonte, M.-B., Leonhardt, A., Borst, A., Mauss, A.S., 2021. Aerial course stabilization is impaired in motion-blind flies. J. Exp. Biol. 224, jeb242219.

Maimon, G., Straw, A.D., Dickinson, M.H., 2010. Active flight increases the gain of visual motion processing in Drosophila. Nat. Neurosci. 13, 393–399.

Mronz, M., Lehmann, F.-O., 2008. The free-flight response of Drosophila to motion of the visual environment. J. Exp. Biol. 211, 2026–2045.

Muijres, F.T., Elzinga, M.J., Iwasaki, N.A., Dickinson, M.H., 2015. Body saccades of Drosophila consist of stereotyped banked turns. J. Exp. Biol. 218, 864–875.

Muijres, F.T., Elzinga, M.J., Melis, J.M., Dickinson, M.H., 2014. Flies evade looming targets by executing rapid visually directed banked turns. Science. 344, 172–177.

Poulet, J.F.A., Hedwig, B., 2006. The Cellular Basis of a Corollary Discharge. Science. 311, 518–522.

Reiser, M.B., Dickinson, M.H., 2008. A modular display system for insect behavioral neuroscience. J. Neurosci. Methods 167, 127–139.

Schilstra, C., Hateren, J.H., 1999. Blowfly flight and optic flow. I. Thorax kinematics and flight dynamics. J. Exp. Biol. 202, 1481–1490.

Schilstra, C., van Hateren, J.H., 1998. Stabilizing gaze in flying blowflies. Nature 395, 654.

Schnell, B., Joesch, M., Forstner, F., Raghu, S. V, Otsuna, H., Ito, K., Borst, A., Reiff, D.F., 2010. Processing of horizontal optic flow in three visual interneurons of the Drosophila brain. J. Neurophysiol. 103, 1646–1657.

Schnell, B., Ros, I.G., Dickinson, M.H., 2017. A Descending Neuron Correlated with the Rapid Steering Maneuvers of Flying Drosophila. Curr. Biol. 27, 1200–1205.

Schnell, B., Weir, P.T., Roth, E., Fairhall, A.L., Dickinson, M.H., 2014. Cellular mechanisms for integral feedback in visually guided behavior. Proc. Natl. Acad. Sci. U. S. A. 111, 5700–5705.

Scott, E.K., Raabe, T., Luo, L., 2002. Structure of the vertical and horizontal system neurons of the lobula plate in Drosophila. J. Comp. Neurol. 454, 470–481.

Sommer, M.A., Wurtz, R.H., 2008. Brain Circuits for the Internal Monitoring of Movements. Annu. Rev. Neurosci. 31, 317–338.

Suver, M.P., Huda, A., Iwasaki, N., Safarik, S., Dickinson, M.H., 2016. An Array of Descending Visual Interneurons Encoding Self-Motion in Drosophila. J. Neurosci. 36, 11768–11780.

Tammero, L.F., Dickinson, M.H., 2002. The influence of visual landscape on the free flight behavior of the fruit fly Drosophila melanogaster. J. Exp. Biol. 205, 327–343.

Tammero, L.F., Dickinson, M.H., 2002. Collision-avoidance and landing responses are mediated by separate pathways in the fruit fly, Drosophila melanogaster. J. Exp. Biol. 205, 2785–2798.

Von Holst, E., Mittelstaedt, H., 1950. The principle of reafference. Naturwissenschaften 37, 464–476.

von Reyn, C.R., Breads, P., Peek, M.Y., Zheng, G.Z., Williamson, W.R., Yee, A.L., Leonardo, A., Card, G.M., 2014. A spike-timing mechanism for action selection. Nat. Neurosci. 17, 962–970.

Wilson, R.I., Laurent, G., 2005. Role of GABAergic Inhibition in Shaping Odor-Evoked Spatiotemporal Patterns in the Drosophila Antennal Lobe. J. Neurosci. 25, 9069–9079.

Wurtz, R.H., 2008. Neuronal mechanisms of visual stability. Vision Res. 48, 2070–2089.

