## Supplementary Figure for "Multiple mechanisms mediate the suppression of motion vision during escape maneuvers in flying *Drosophila*"

### SUPPLEMENTARY MATERIAL

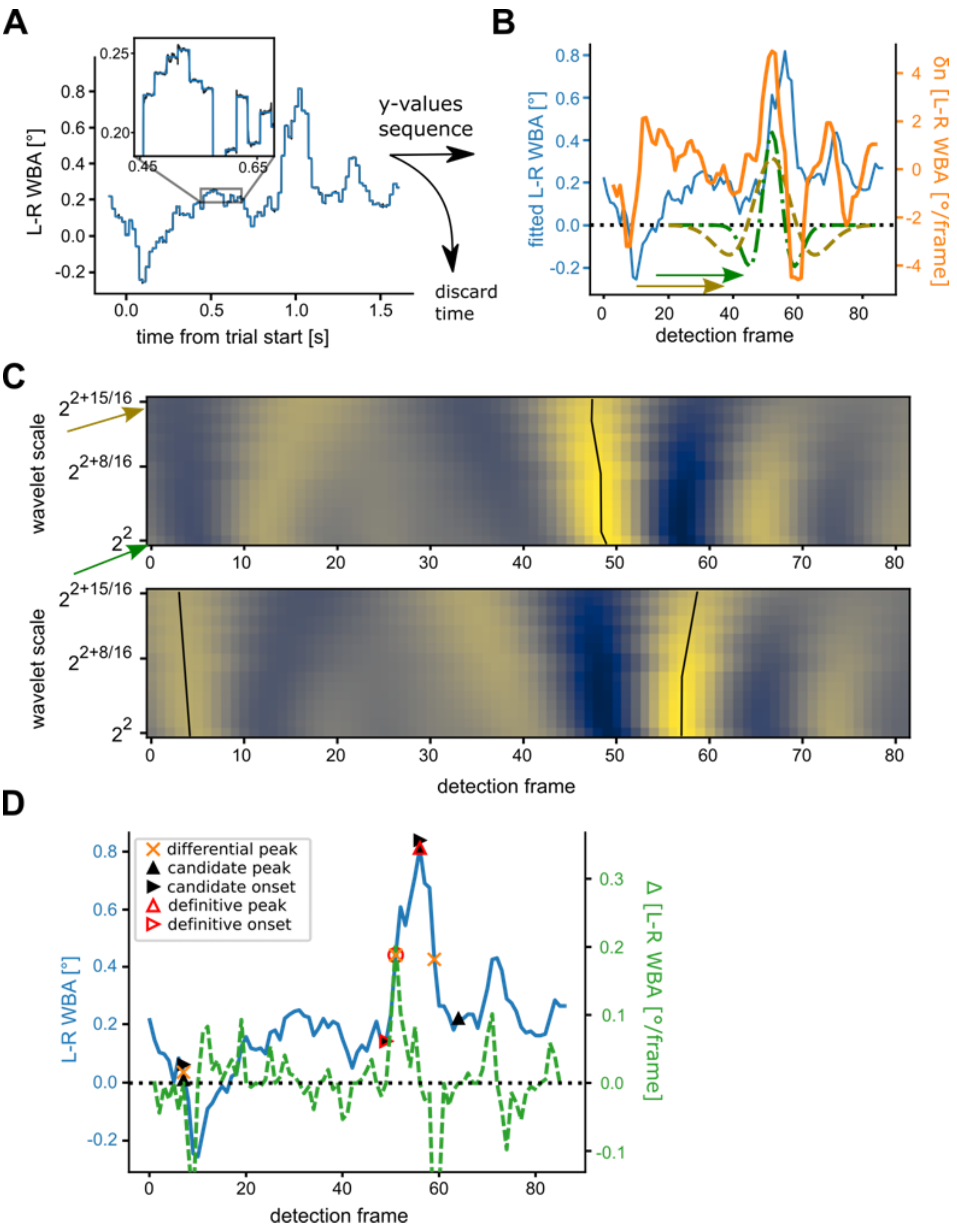


**Supplementary Figure 1**

Illustration of the saccade detection algorithm. (**A**) The wingbeat amplitude signal is corrupted by additive noise, which persists in the L-R WSA (black). Using the prior knowledge of approximate camera frame rate, a piece-wise-constant function is fitted to it (blue), to retrieve a distortion-free downsampled signal as a sequence of y-values. (**B**) Y-values form a continuous-value time-series (blue), which is differentiated using the central difference theorem (orange). Wavelet functions of the Ricker (derivative of a gaussian) family are applied as filters for peaks in the derived signal with scales between 4 (green) and 8 (brown) similar to a convolution. (**C**) Exemplary result of continuous wavelet transform (CWT) in the given range. Top shows CWT of positive signal, bottom is CWT of negative, arrows point at scales of wavelets in (B). Ridges across wavelet scales correspond to sharp transients, which are candidates for saccades. Rightward saccades correspond to peaks in the positive signal, and leftward saccades to peaks in the negative. (**D**) Ridge locations in CWT in positive and negative signals are treated as candidate peaks and additional post-selection criteria used to identify sufficiently prominent, stand-alone peaks in L-R WSA as saccades. Saccade onset and peak times are identified as time points closest to zero in the simple (backward difference) derivative.
